# Trends in female lifespan in South Korea, 1987–2016

**DOI:** 10.1101/278291

**Authors:** Byung Mook Weon

## Abstract

South Korea shows a remarkable rapid increase in lifespan in recent decades. Employing a mathematical model that is appropriate for human survival curves, we evaluate current trends in female lifespan for South Korea over three recent decades, 1987–2016, and predict coming trends in female lifespan until 2030. From comparative analyses with industrialized countries such as Japan, France, Australia, Switzerland, UK, Sweden, and USA, we confirm that South Korea has the highest increase rate of female lifespan in recent decades, and estimate that maximum lifespan would reach 125 years and characteristic life would surpass 95 years for South Korean female by 2030. South Korea would deserve much attention in study on human health and longevity as the longest-lived country in coming decades.

## Introduction

Statistical analysis of human lifespan is a central topic in biogerontology. Life expectancy of humans has risen steadily and rapidly over the past 150 years in most countries and keeps increasing in the 21th century (Wilmoth et al. 2000; Oeppen and Vaupel 2006; Olshansky 2016). A recent study for 35 industrialized countries shows a reliable projection: there is more than 50% probability that by 2030, female life expectancy is likely to break the 90-year barrier (Kontis et al. 2017). Most importantly, South Korea is predicted to have the highest female life expectancy at birth by 2030 (Kontis et al. 2017). This prediction is achieved by applying a mathematical ensemble of 21 forecasting models to reliable statistics data about life tables for humans, which would be important in planning for health and social services and pensions (Kontis et al. 2017).

The precise mathematical modeling of human survival curves is essential in evaluating and predicting current and future trends in human lifespan, which is a fundamental task in human aging and demography research (Weon 2015; Petrascheck and Miller 2017; Ruby et al. 2018; Beltrán-Sánchez and Finch 2018). So far, many mathematical models for human survival curves have been proposed, including the Gompertz, Weibull, Heligman-Pollard, Kannisto, quadratic, and logistic models (see references in Weon 2015). In the recent work, we have suggested a useful mathematical model that is appropriate to depict human survival curves with complexity in shape by modifying a stretched exponential (equivalently Kohlrausch-Williams-Watts or Weibull) function (Weon and Je 2009; 2011; 2012; Weon 2015). In the present work, by incorporating this methodology to the reliable life table datasets taken from the Korean Statistical Information Service (KOSIS, http://kosis.kr/eng), we evaluate current trends in female lifespan for South Korea over three recent decades, from 1987 to 2016, and estimate coming trends in female lifespan until 2030. This analysis for South Korea is compared with that for Japan, France, Australia, Switzerland, UK, Sweden, and USA, which are the leading countries in longevity (Kontis et al. 2017).

The survival curve is described as the survival rate, s(x), as a function of age, x. Mathematically the mortality rate (equivalently the hazard function or the force of mortality), μ(x) = −dln(s(x))/dx, is computed from the derivative of the survival rate with respect to age. The important feature of s(x) is that s(x) always monotonically decreases with age from 1 to 0. To describe the human survival curves, a modified stretched exponential function has been developed as a form of s(x) = exp(–(x/α)^β(x)^) where the stretched exponent, β(x), is defined as β(x) = ln[–ln(s(x))]/ln(x/α) and the characteristic life, α, is given at s(α) = exp(–1) that is practically obtained at the interception point between s(x) and s(α) = exp(–1) (Weon and Je 2009; 2011; 2012; Weon 2015). The characteristic life is a good alternative to the life expectancy at birth, ε (Wrycza and Baudisch 2014; Weon and Je 2012). The age dependence of the stretched exponent is the critical feature of the modified stretched exponential function, which is fundamentally different from the classical stretched exponential (equivalently the Kohlrausch-Williams-Watts (Kohlrausch 1854; Williams and Watts 1970) or the Weibull (Weibull 1951)) function (Weon and Je 2009; 2011; 2012; Weon 2015). The characteristic life and the stretched exponent are useful to respectively describe the scale effect (associated with ‘living longer‘, characterized by α) and the shape effect (associated with ‘growing older‘, characterized by β(x)) of the individual survival curve (Weon and Je 2012). To describe the β(x) patterns at very old ages, the quadratic formula of β(x) = β_0_ + β_1_x + β_2_x^2^ is applicable (Weon and Je 2009; Weon 2015). The quadratic patterns in β(x) certainly lead to the occurrence of the (mathematical) maximum lifespan, ω, which is obtained at the age for β(x) = – xln(x/α) dβ(x)/dx by the constraint of ds(x)/dx → 0 (Weon and Je 2009). This methodology is beneficial in modeling the human mortality curves in very old age (Weon 2015).

In this study, we evaluate the survival curves for South Korean female in the most recent decades, 1987–2016, taken from the Korean Statistical Information Service (KOSIS, http://kosis.kr/eng), by adopting the modified stretched exponential function. Eventually, we demonstrate the current and future trends (through evaluation for 1987–2016 and prediction by 2030) in the maximum lifespan and the characteristic life for the female survival curves in South Korea.

## Analysis

Figure 1 shows how we evaluate the survival curves for South Korean female in the most recent decades, 1987–2016. The survival curves for women were obtained at the complete life tables taken from the Korean Statistical Information Service (KOSIS, http://kosis.kr/eng) [accessed on 10 February 2018]. The summarized survival curves in Fig. 1a show the historic trends in the survival rates for South Korean female over three recent decades (1987–2016). The characteristic life, α, indicating the scale effect (associated with ‘living longer‘) of the individual survival curve, was measured for each survival curve through the graphical analysis at s(α) = exp(-1) ≈ 0.367879 and the historic trends in the α values were summarized in Fig. 1b. The age-dependent stretched exponent, β(x), indicating the shape effect (associated with ‘growing older‘) of the individual survival curve, was obtained from β(x) = ln[-ln(s(x))]/ln(x/α) as shown in Fig. 1c. Particularly as demonstrated in Fig. 1d, the old-age patterns of β(x) at ages over 60 years could be fitted with the quadratic formula as β(x) = β_0_ + β_1_x + β_2_x^2^ (Weon and Je 2009; 2011; Weon 2015).

**Figure 1.**
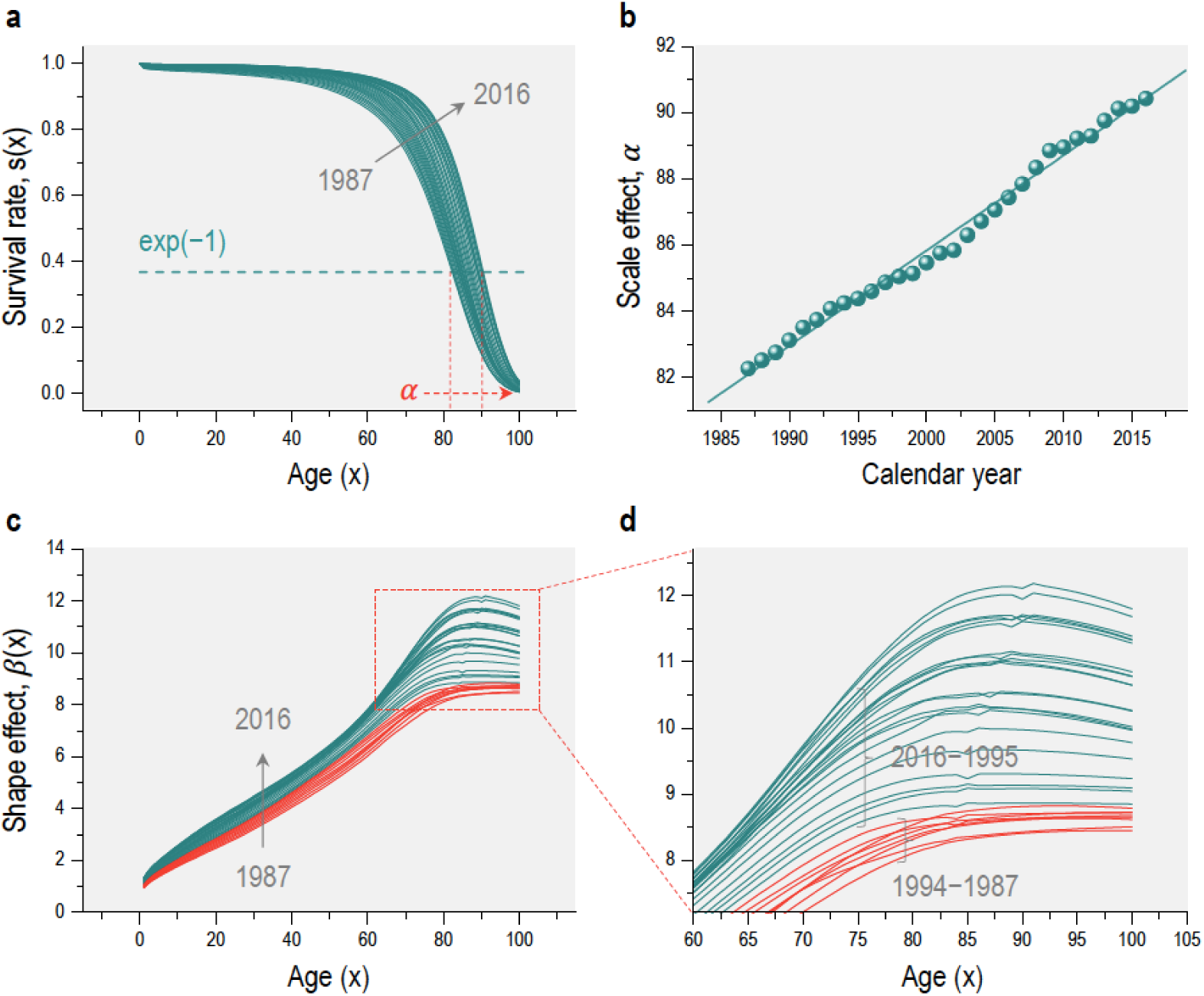
Historic trends in survival curves for South Korean female in recent decades, 1987–2016. (a) The survival curves were obtained at the complete life tables taken from the Korean Statistical Information Service (KOSIS, http://kosis.kr/eng) [accessed on 10 February 2018]. (b) The characteristic life of the given survival curve was measured for each survival curve through the graphical analysis at s(α) = exp(-1) ≈ 0.367879. (c) The age-dependent stretched exponent of the given survival curve was obtained by computing β(x) = ln[-ln(s(x))]/ln(x/α). (d) The old-age patterns of β(x) at ages over 60 years could be fitted with the quadratic formula of age. The phase transition in β(x) with age, whether β(x) increases with age at x < α or decreases with age at x > α, appears for 1995–2016 but not for 1987–1994.

The estimated maximum lifespan, ω, and estimated characteristic life, α, from 2002 to 2030 are summarized in Fig. 2. The maximum lifespan was determined by the mathematical constraint of ds(x)/dx → 0 (Weon and Je 2012; Weon 2015). The fitting of β(x) with the quadratic formula enables us to measure the estimate of the maximum lifespan at the specific age of γ(x) = –xln(x/α) dβ(x)/dx (that is graphically measurable at the points β(x) = γ(x)) (Weon 2015). This estimation is practically available if and only if the phase transition exists in the old-age β(x) pattern. Therefore, we estimate the ω values after the calendar year of 2002 (as marked by the stars in Fig. 2).

**Figure 2.**
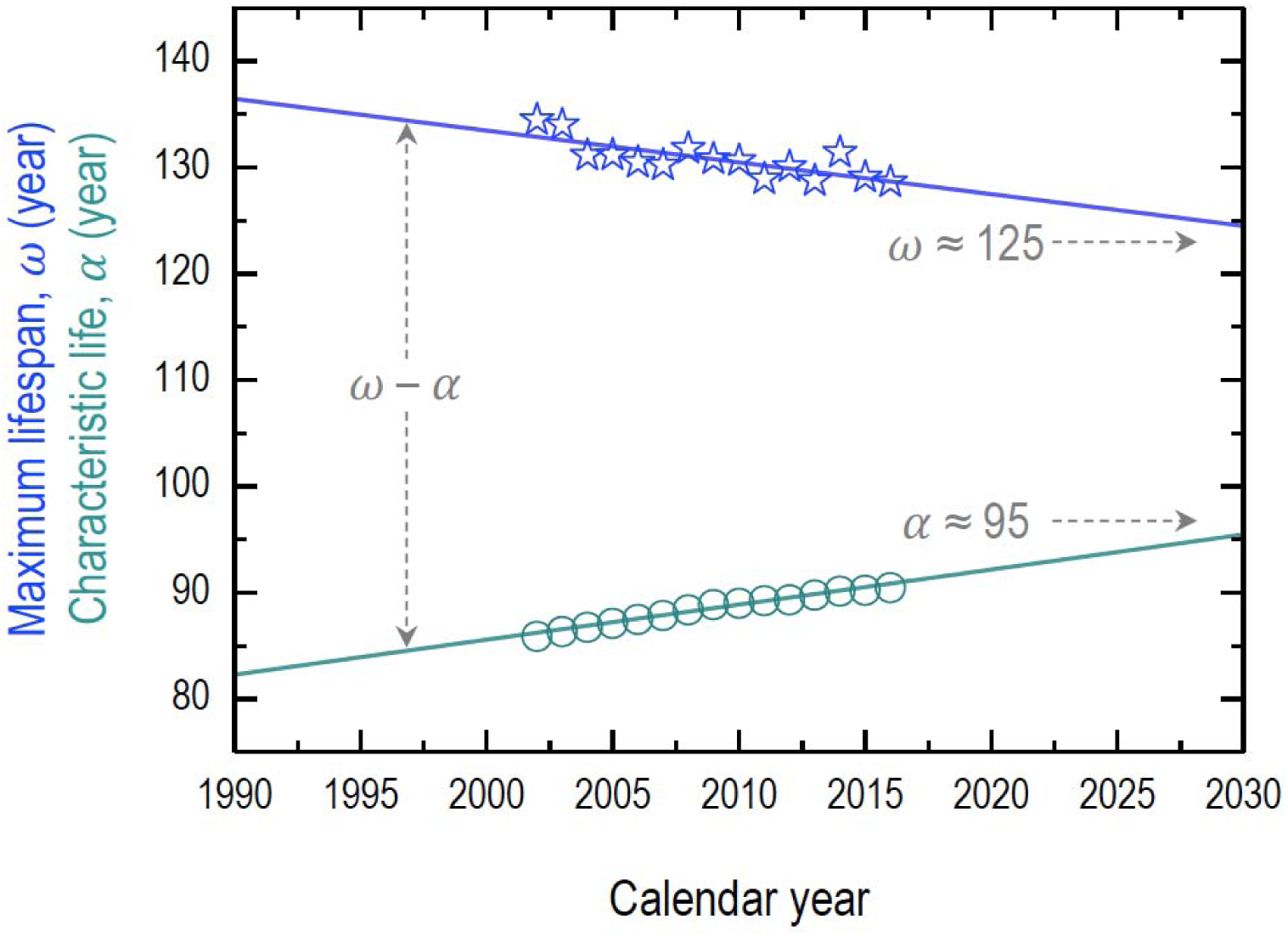
Current and future trends of the maximum lifespan, ω, and the characteristic life, α, for South Korean female. The maximum lifespan gradually decreases at a constant rate of ∼2.99 years per decade, while the characteristic life gradually increases at a constant rate of ∼3.30 years per decade. The maximum lifespan would be expected to reach ∼125 years by 2030 and the characteristic life would surpass ∼95 years by 2030.

## Results and Discussion

The term ‘lifespan’ describes how long an individual can live and the (observed) maximum lifespan is the age reached by the longest-lived member of a species, while the life expectancy is a population-based estimate of expected duration of life for individuals at any age, based on a statistical life table (Olshansky 2016). Our methodology enables us to estimate the characteristic life, α, and the (mathematical) maximum lifespan, ω, as good measures of the human lifespan.

The survival curves for South Korean female, as illustrated in Fig. 1a, show that s(x) gradually decreases with age and historically shifts rightwards and towards a rectangular shape over three recent decades, 1987–2016, as consistent with Swedish female (Weon and Je 2012; Weon 2015). The characteristic life for South Korean female was depicted as a function of the calendar year (1987–2016) in Fig. 1b, showing that the α value linearly increases at a constant rate of ∼2.88 years per decade (the solid line in Fig. 1b). This steady increase rate for South Korea is much higher than the rate of ∼1.2 years per decade for Swedish female (Weon and Je 2012; Weon 2015). The historic trends in the characteristic life, representing the scale effect (associated with ‘living longer‘), clearly show that the female lifespan in South Korea has been living longer during 1987–2016.

The modified stretched exponential function was successfully applicable to the survival curves and to the age-dependent stretched exponents, β(x), (marked by the shape effect) in Figs. 1c & 1d., respectively for all ages (Fig. 1c) and for old ages over 60 years (Fig. 1d). Most remarkably, the β(x) curve smoothly varies with age at old ages over 60 years and the old-age β(x) patterns over 90 years can be depicted with the quadratic function of age.

Typically, the β(x) patterns for long-lived populations show the phase transition with age: β(x) increases with age at x < α and decreases with age at x > α (Weon and Je 2009). The phase transition appears in South Korean female datasets after the calendar year of 1995: particularly we find phase transitions for 1995–2016 and no transitions for 1987–1994, as separately marked in Fig. 1d. Considering the old-age β(x) patterns, the female lifespan of South Korea would experience a critical change before and after the calendar year of 1995. The historic trends in the age-dependence in β(x), representing the shape effect (associated with ‘growing older‘), clearly show how the female lifespan in South Korea has been growing older during 1987–2016.

The current and future trends of the maximum lifespan, ω, and the characteristic life, α, are summarized in Fig. 2. Here, it is clarified that for the South Korean female datasets during 2002–2016, the maximum lifespan gradually decreases at a constant rate of ∼2.99 years per decade, while the characteristic life gradually increases at a constant rate of ∼3.30 years per decade. Assuming continuity in the increments, the estimates of ω and α suggest that the maximum lifespan would reach ∼125 years and the characteristic life would surpass ∼95 years by 2030. The change rates for South Korean female are much higher than those of ∼1.6 years per decade in the ω values and ∼1.2 years per decade in the α values for Swedish female (Weon 2015). The estimates of the maximum lifespan and the characteristic life eventually become closer together over time: this tendency for South Korean female is consistent with Swedish female (Weon 2015). This result indicates that the survival curves have been increasingly concentrated at old ages, demonstrating the population aging (Anderson and Hussey 2000; Robine and Michel 2004). As the characteristic life becomes closer to the maximum lifespan (Fig. 2), the survival curves become increasingly concentrated at very old ages, corresponding to the retangularization of survival curves (Fries 1980; Weon and Je 2011). This tendency is relevant to the compression of morbidity, which becomes more important in very old age and would be responsible for the concentration of the very old populations (Weon 2015).

The detailed comparison in the increase rate of the characteristic life, α, for South Korea with that for major industrialized countries is shown in Fig. 3. The increase rate in α was obtained from Fig. 1b for South Korea, 1987–2016, and from literature (Weon and Je 2012) for other countries, 1980–2010 (the scale bars taken from the standard errors by linear fits). The increase rate in α per decade was evaluated as ∼2.88 years for South Korea, ∼2.71 years for Japan, ∼1.96 years for France, ∼1.85 years for Australia, ∼1.61 years for Switzerland, ∼1.49 years for UK, ∼1.24 years for Sweden, and ∼0.76 years for USA. This analysis confirms the highest increase rate of the characteristic life for women in South Korea among the major high-income countries including Japan, France, Australia, Switzerland, UK, Sweden, and USA.

**Figure 3.**
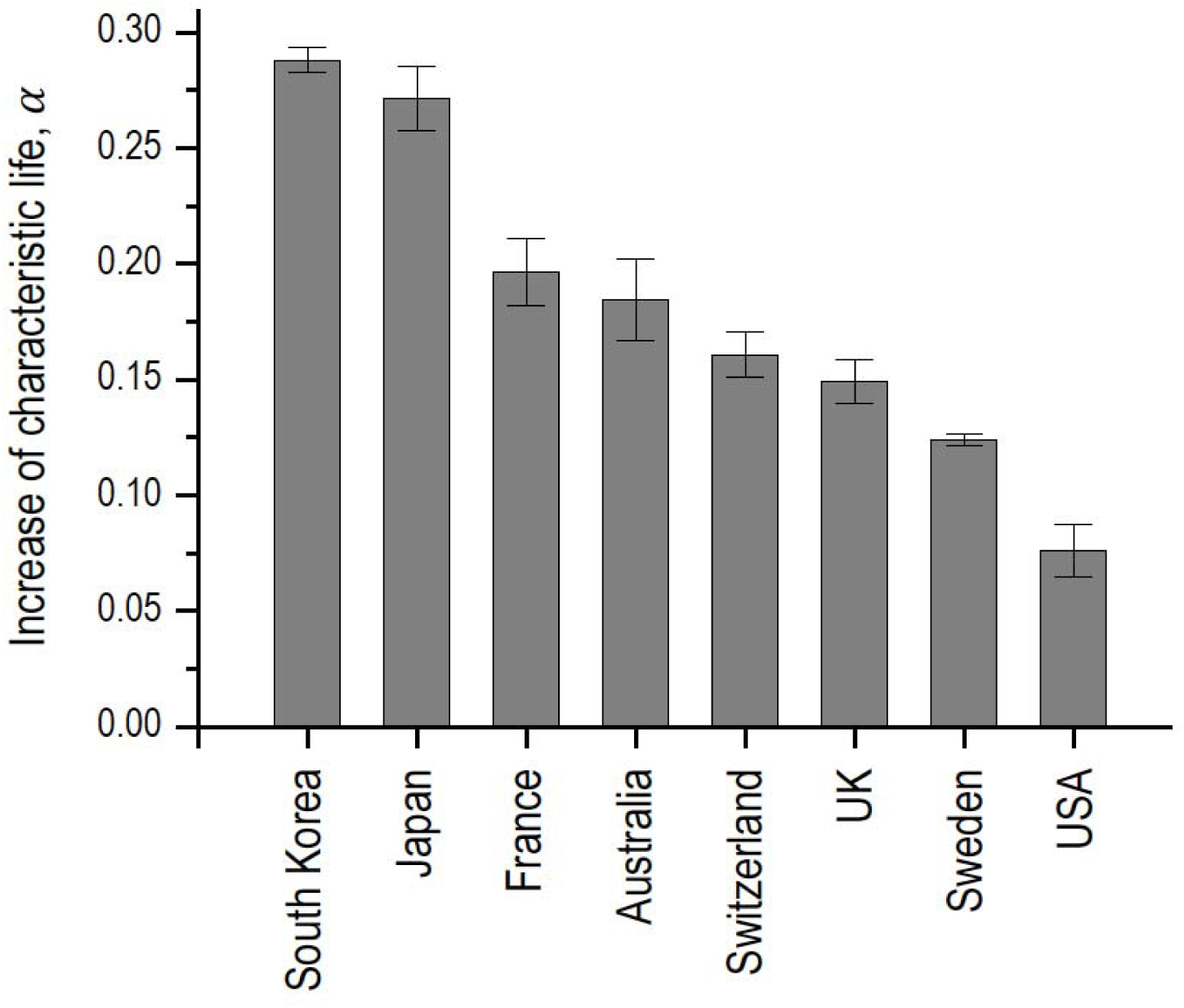
Comparison in the increase rate of the characteristic life, α, for South Korea with that for major industrialized countries. The increase rate of the α values per decade was evaluated as ∼2.88 years for South Korea, ∼2.71 years for Japan, ∼1.96 years for France, ∼1.85 years for Australia, ∼1.61 years for Switzerland, ∼1.49 years for UK, ∼1.24 years for Sweden, and ∼0.76 years for USA.

To demonstrate the association of the characteristic life, α, with the life expectancy at birth, ε, we correlate the annual increase rates of the α and the ε values in Fig. 4. The ε values for 1980–2015 were taken from the Organization for Economic Cooperation and Development (OECD 2017) and the increase rates of the ε values were achieved by linear fits (the scale bars taken from the standard errors). The increase rates of the ε values per decade were evaluated as ∼4.15 years for South Korea, ∼2.33 years for Japan, ∼2.15 years for France, ∼1.96 years for Australia, ∼1.78 years for Switzerland, ∼1.93 years for UK, ∼1.43 years for Sweden, and ∼1.06 years for USA. The annual increase rate of the ε values (marked by ‘m‘) is proportionally correlated to the annual increase rate of the α values (marked by ‘n‘) as n ≈ 1.173m – 0.046 (adj. R^2^ ∼ 0.85243) except for South Korea. Interestingly, the annual increase rate of the ε values for South Korea looks vastly superior to that for other countries.

**Figure 4.**
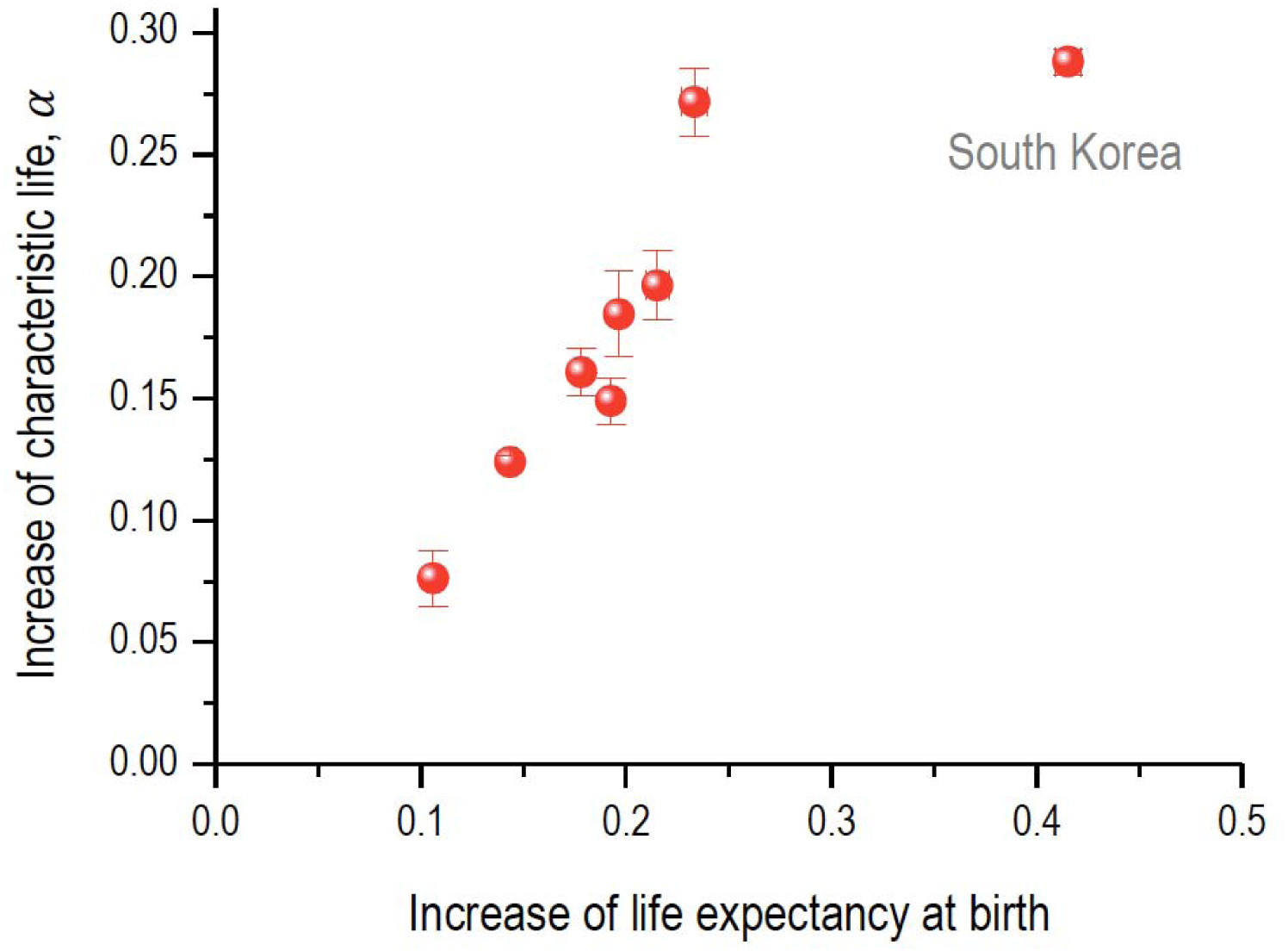
Association of the characteristic life, α, with the life expectancy at birth, ε, for South Korea and major industrialized countries. The increase rates of the ε values per decade were evaluated as ∼4.15 years for South Korea, ∼2.33 years for Japan, ∼2.15 years for France, ∼1.96 years for Australia, ∼1.78 years for Switzerland, ∼1.93 years for UK, ∼1.43 years for Sweden, and ∼1.06 years for USA.

The highest increase rate in the female lifespan (in both α and ε) for South Korea in Fig. 4 can explain why the female life expectancy at birth in South Korea is sixth ranked in 2010 but expected to be first ranked in 2030 among the 35 industrialized countries (Kontis et al. 2017). So far, little attention has been given to South Korea in research on human health and longevity except for few recent reports (Kontis et al. 2017). Evidently, South Korea has the highest increase rate in the female lifespan in the world.

We discuss how South Korea has acquired the highest increase rate in the female lifespan. The recent increase in the numbers of older people in developed countries is a consequence of advances in hygiene and biomedicine as well as an artefact of human civilization (Hayflick 2000). South Korea has experienced rapid economic development and a substantial increase in life expectancy in an extremely short period, between 1970 and 2010, mainly due to the rapid processes of industrialization and urbanization (Kim et al. 1996; Bahk et al. 2017). South Korea is projected to become the first country where the life expectancy at birth will exceed 90 years for women (Khang et al. 2016). The recent industrialization and the subsequent improvements in living standards, nutrition, and health care have often been cited as major contributions to the remarkable improvements in health for South Korea (Yang et al. 2010). The rapid increases in life expectancy in South Korea could be mostly achieved by reductions in infant mortality and in decreases related to infections and blood pressure (Yang et al. 2010). More recent gains in South Korea have been largely due to postponement of death from chronic diseases (Yang et al. 2010). These gains were mainly due to broad-based inclusive improvements in economic status and social capital (including education), which improved childhood and adolescent nutrition (NCD-RisC 2016a), expanded access to primary and secondary health care, and facilitated rapid scale-up of new medical technologies (Yang et al. 2010). South Korea has also maintained lower body-mass index and blood pressure than other industrialized countries (NCD-RisC 2016b; 2017), and lower smoking in women. Additionally, South Korea might have lower health inequalities (for cancer and cardiovascular disease mortality, and for self-reported health status) than some of their western counterparts, especially for women (Di Cesare et al. 2013; OECD 2015; Khang et al. 2016). The death rates from all causes in South Korea decreased significantly in both genders in the last three decades except for a period following the economic crisis in the late 1990s (Lim et al. 2014). Looking at the trends in infectious disease mortality for South Korea, 1983–2015, infant mortality caused by infectious diseases has substantially decreased, while death rates from infectious disease for elderly populations with lower education levels and subgroups susceptible to respiratory infections and sepsis has not decreased overall (Choe et al. 2018).

The lifespan trends for South Korean female may give a clue for the fundamental question about how human aging evolves or whether there is a limit to human lifespan (Carnes and Witten 2014; Dong et al. 2016; Olshansky 2016; Newman and Easteal 2017; Marck et al. 2017). The heterogeneity in population and the dynamics of heterogeneous populations would become important in interpreting human lifespan (Carnes and Olshansky 2001; Vaupel 2010). Questions about whether the upper limit to the maximum lifespan is fixed or flexible have been topics of continuous debate (Dong et al. 2016; Ben-Haim et al. 2017; Brown et al. 2017; de Beer 2017; Hughes and Hekimi 2017; Lenart and Vaupel 2017; Rozing et al. 2017). The precise mathematical evaluation of the survival curves would be essential in achieving reliability and reproducibility in aging research (Petrascheck and Miller 2017; Ruby et al. 2018; Beltrán-Sánchez and Finch 2018) as well as in demonstrating the effectiveness of anti-aging therapies to slow the aging process (de Magalhães 2017). Our mathematical model is quite appropriate to evaluate the human survival curves with heterogeneity and complexity in shape, as demonstrated herein.

A steady increase in lifespan has been recorded in recent decades for most industrialized countries. This study shows evidence for that South Korea has a remarkable rapid increase in lifespan among industrialized countries. By adopting the modified stretched exponential survival function, we measure the increase rate of female lifespan for South Korea over three recent decades, from 1987 to 2016, and predict the upcoming trends in female lifespan until 2030. From comparative analyses with Japan, France, Australia, Switzerland, UK, Sweden, and USA, we confirm that South Korea has the highest increase rate of female lifespan over three recent decades, and estimate that the maximum lifespan would reach 125 years and the characteristic life would surpass 95 years for women by 2030. This result implies that South Korea would be the longest-lived country in coming decades and would provide many hints to human health and longevity. South Korean female datasets can be a representative population dataset for further study on the dynamics of human longevity. Our methodology would contribute to the better tracking and forecasting of human aging statistics, which would be of great importance for future population and policy.

## Acknowledgements

This research was supported by Basic Science Research Program through the National Research Foundation of Korea (NRF) funded by the Ministry of Education (NRF-2016R1D1A1B01007133).

